# Identification of differentially expressed genes in hepatocellular carcinoma by integrated bioinformatic analysis

**DOI:** 10.1101/570846

**Authors:** Guangxin Yan, Zhaoyu Liu

**Affiliations:** Department of Radiology, Shengjing Hospital affiliated to China Medical University, Shenyang, China

**Keywords:** HCC, GEO, TCGA, DEGs, bioinformatic analysis

## Abstract

Hepatocellular carcinoma is one of the most common tumors in the world and has a high mortality rate. This study elucidates the mechanism of hepatocellular carcinoma- (HCC) related development. The HCC gene expression profile (GSE54238, GSE84004) was downloaded from Gene Expression Omnibus for comprehensive analysis. A total of 359 genes were identified, of which 195 were upregulated and 164 were downregulated. Analysis of the condensed results showed that “extracellular allotrope” is a substantially enriched term. “Cell cycle”, “metabolic pathway” and “DNA replication” are three significantly enriched Kyoto Encyclopedia of Genes and Genomespathways. Subsequently, a protein-protein interaction network was constructed. The most important module in the protein-protein interaction network was selected for path enrichment analysis. The results showed that *CCNA2, PLK1, CDC20, UBE2C* and *AURKA* were identified as central genes, and the expression of these five hub genes in liver cancer was significantly increased in The Cancer Genome Atlas. Univariate regression analysis was also performed to show that the overall survival and disease-free survival of patients in the high expression group were longer than in the expression group. In addition, genes in important modules are mainly involved in “cell cycle”, “DNA replication” and “oocyte meiosis” signaling pathways. Finally, through upstream miRNA analysis, mir-300 and mir-381-3p were found to coregulate *CCNA2*, *AURKA* and *UBE2C*. These results provide a set of targets that can help researchers to further elucidate the underlying mechanism of liver cancer.

## 1. Introduction

Liver cancer is one of the most common cancers in the world, and mainly includes three different pathological types comprising hepatocellular carcinoma (HCC), intrahepatic bile duct (ICC) and HCC-ICC hybrid types, among which HCC accounts for 85% to 90% of all HCCs. The incidence and mortality of HCC increase with age, and the incidence of HCC is about three times higher in men than in women[1]. Although HCC treatment has made great progress in recent years, the 5-year survival rate of HCC patients is still <25%[2]. Therefore, more research are needed to understand the molecular mechanisms of HCC development and progression, which is important to develop more effective diagnostics and treatments.

At present, gene microarray technology has been widely used to collect gene chip expression profiling data and to study gene expression profiles in many human cancers. A large amount of data has been published in public database platforms, and researchers have integrated these databases to find valuable information about cancer pathogenesis. Lau et al. identified approximately 4,000 genes in 10 pairs of HCC and non-tumor tissues by microarray technology[3]. Zhou et al. analyzed the mRNA expression profiles of a large number of human HCC specimens from four independent Gene Expression Omnibus(GEO) microarray datasets and identified key genes and potential molecular mechanisms for HCC development[4]. Additionally, Wang et al. provided a pipeline for the identification of prognostic signatures for HCC OS prediction[5]. Microarray technology has since been applied to the study of genetic changes in HCC, has defined several different genetic variants of the disease, and has identified genetic features that predict poor outcomes and metastasis [6–8]. Although the cellular and molecular genetic alterations of HCC have been more well-understood through these technologies, the molecular mechanisms have not yet been fully elucidated.

To further investigate the molecular mechanisms of HCC, we downloaded two microarray datasets, GSE54238 and GSE84004, from GEO to identify differentially expressed genes (DEGs) in HCC and normal liver tissues. In addition, we integrated bioinformatics into DEG’s Gene Ontology (GO) and Kyoto Encyclopedia of Genes and Genomes (KEGG) analysis, then constructed a protein-protein interaction (PPI) network and analyzed the central genes. We then downloaded relevant data from The Cancer Genome Atlas (TCGA) to verify and evaluate the key genes. In conclusion, this study will help to better understand the occurrence and development of HCC and will also contribute to the early diagnosis and treatment of HCC.

## 2. Materials and methods

### 2.1 Gene-expression data

Data were downloaded from the Gene Expression Omnibus (GEO, http://www.ncbi.nlm.nih.gov/geo), a public repository for data storage. Two RNA expression datasets of HCC GSE54238 and GSE84004 were used in this study. The dataset GSE54238, which is based on GPL16955(Arraystar human lncRNA microarray V1 100309), included ten normal liver, ten chronic inflammatory liver, ten cirrhotic liver, thirteen early HCC and thirteen advanced HCC samples; the dataset GSE84004 based on GPL22109 (NimbleGenHuman 100309AShuman100426pz array) included thirty eight non-tumor liver, and thirty eight liver cancer samples.

### 2.2 DEG analysis

Raw data were converted into an expression matrix, which was subsequently normalized with the robust multi-array average algorithm in the Affy packagein R (https://www.r-project.org/). The *t*-test method in the limma R package was subsequently used to identify DEGs between the tumor tissues and adjacent non-tumorous tissue samples. A |log_2_-fold change|>1 and P<0.05 were considered as the threshold values for DEG identification.

### 2.3 GO and KEGG analyses

The Database for Annotation, Visualization and Integrated Discovery (DAVID; http://david.abcc.ncifcrf.gov/)[9] is an essential resource for high-throughput gene analysis that provides researchers with a comprehensive set of functional annotation tools to understand the biological implications behind many genes. GO[10] was performed on the identified DEG using the DAVID database, including molecular function, biological process, cellular components, and KEGG[11] pathway enrichment analysis.

### 2.4 PPI network construction and module analysis

A search tool for identifying interacting genes (STRING, http://string.embl.de/)[12] is a biological database for predicting PPI information. Interactions with a combined score >0.4 were defined as statistically significant. Cytoscape software (version 3.6.1) was used to visualize the integrated regulatory networks. According to the degree levels in the Cytoscape plugin cytoHubba (version 0.1), the top five ranked genes were defined as hub genes. Molecular complex detection (MCODE) was used to screen modules for PPI networks. The standard settings are as follows: degree cutoff = 2, node score cutoff = 0.2, k-core = 2, maximum depth = 100. In addition, the function and path enrichment analysis of the DEGs in the module is performed.

### 2.5 Validation of the expression of hub genes in TCGA

The Cancer Genome Atlas (TCGA, https://cancergenome.nih.gov/)[13]is a cancer database that contains clinical, genomic variation, mRNA expression, miRNA expression, and methylation data, etc., for various human cancers. We obtained clinical information and RNA sequencing data for HCC from TCGA. The prognosis of the five hub genes was assessed using univariate Cox regression analysis and Kaplan-Meier curves.

### 2.6 Upstream miRNA analysis

Based on the information obtained on the miRNA-mRNA pairs in starBase (version 2.0), the upstream mRNAs were respectively predicted. The shared upstream miRNAs of the mRNAs were then identified according to their expression levels.

## 3. Results

### 3.1 DEG analysis

The HCC expression microarray data sets GSE54238 and GSE84004 were normalized. When the GSE54283 data set was screened by the limma package (corrected P value <0.05, logFC > 2), 1690 DEGs were obtained. One thousand and six DEGs were screened from the GSE84004 data set. An expression heat map (Fig. 1) and a volcano plot of the identified DEGs were constructed. Further Venn analysis showed that 359 DEGs were obtained in both data sets, including 195 upregulated and 164 downregulated genes (Fig. 2). The top 10 most significantly up- or downregulated differentially expressed genes are listed in(Table 1).

**Fig 1.**
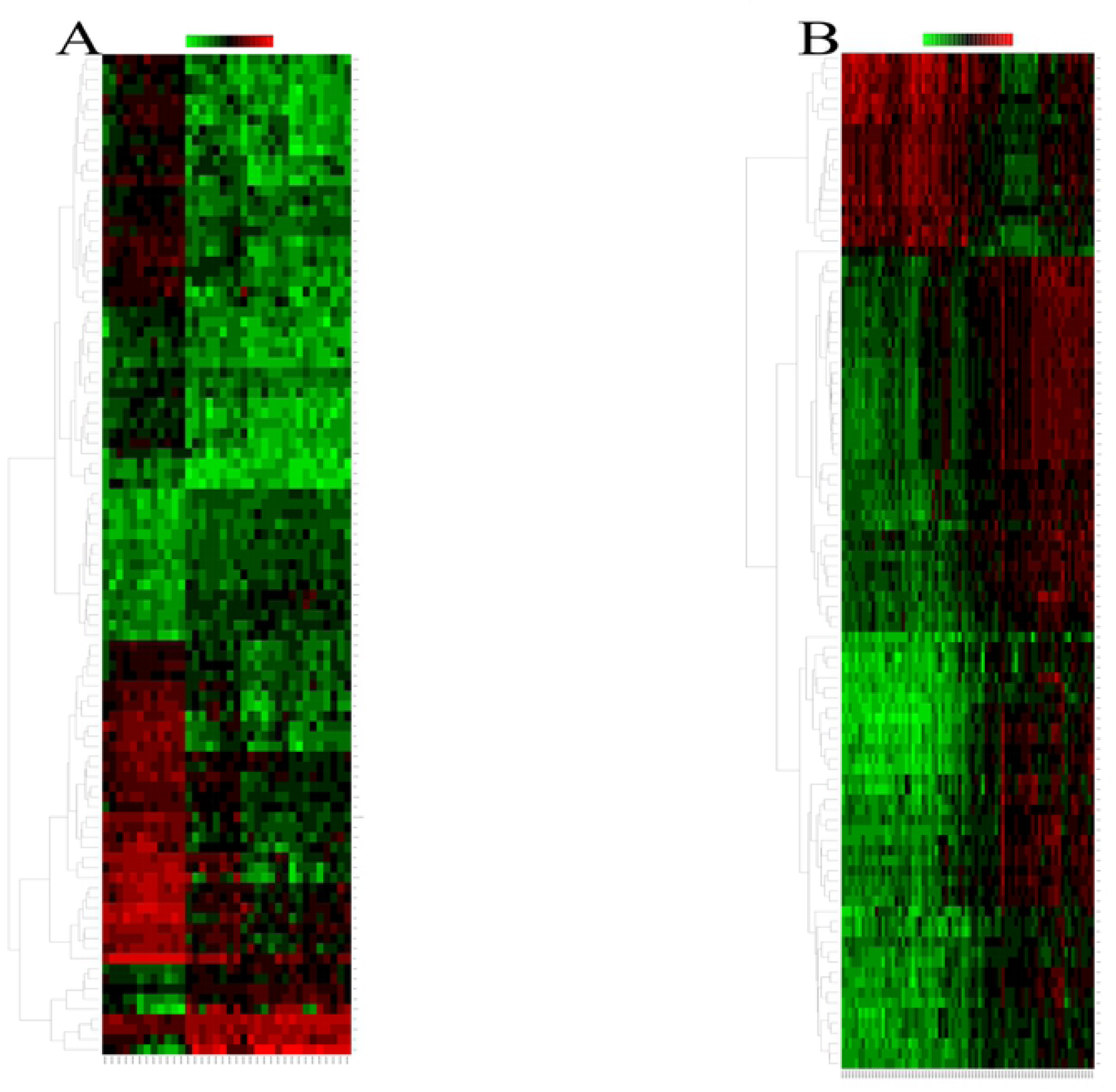
Hierarchical clustering heatmap of the top 100 DEGs. (A) GSE54238 data (B) GSE84004 data. Red indicates upregulated relative gene expression, green indicates downregulated relative gene expression, and black indicates no significant changes in gene expression. DEGs, differentially expressed genes.

**Fig 2.**
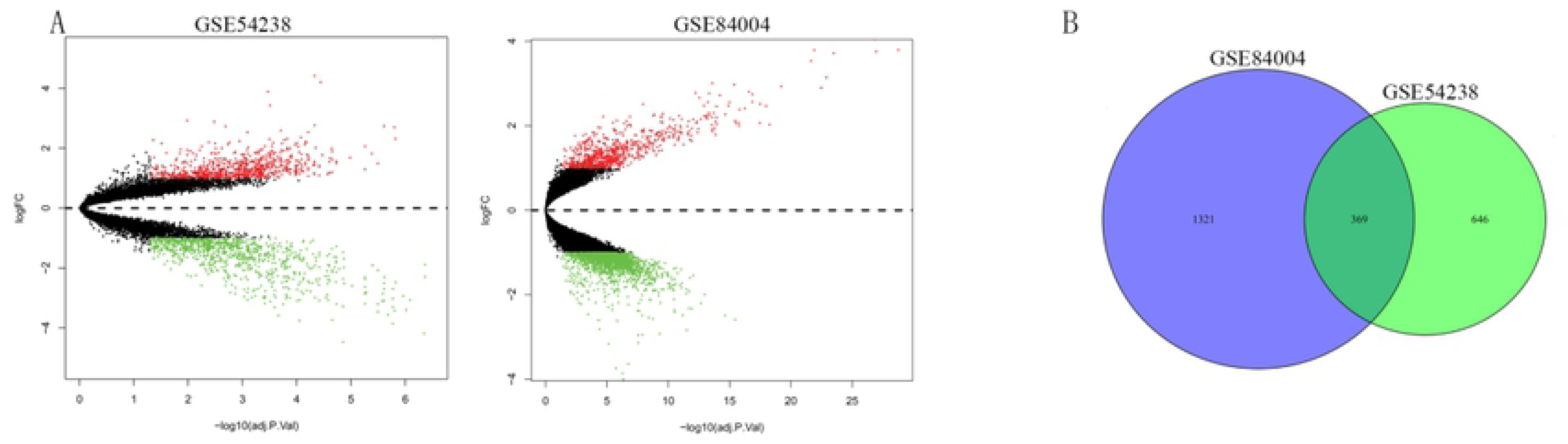
(A) Differential expression of data between two sets of samples. The left is the GSE54238 data and the right is the GSE84004 dataset. The red points represent upregulated genes screened based on |fold change| >1.0 and a corrected P-value of <0.05. The green points represent downregulation of the expression of genes screened based on |fold change| >1.0 and a corrected P-value of <0.05. The black points represent genes with no significant differences in gene expression. (B)Venn diagram illustrating the number of DEGs in GSE54238 and GSE84004 data. Gray intersections represent the common DEGs between two data sets. DEG, differentially expressed gene; FC, fold change.

**Table 1.**
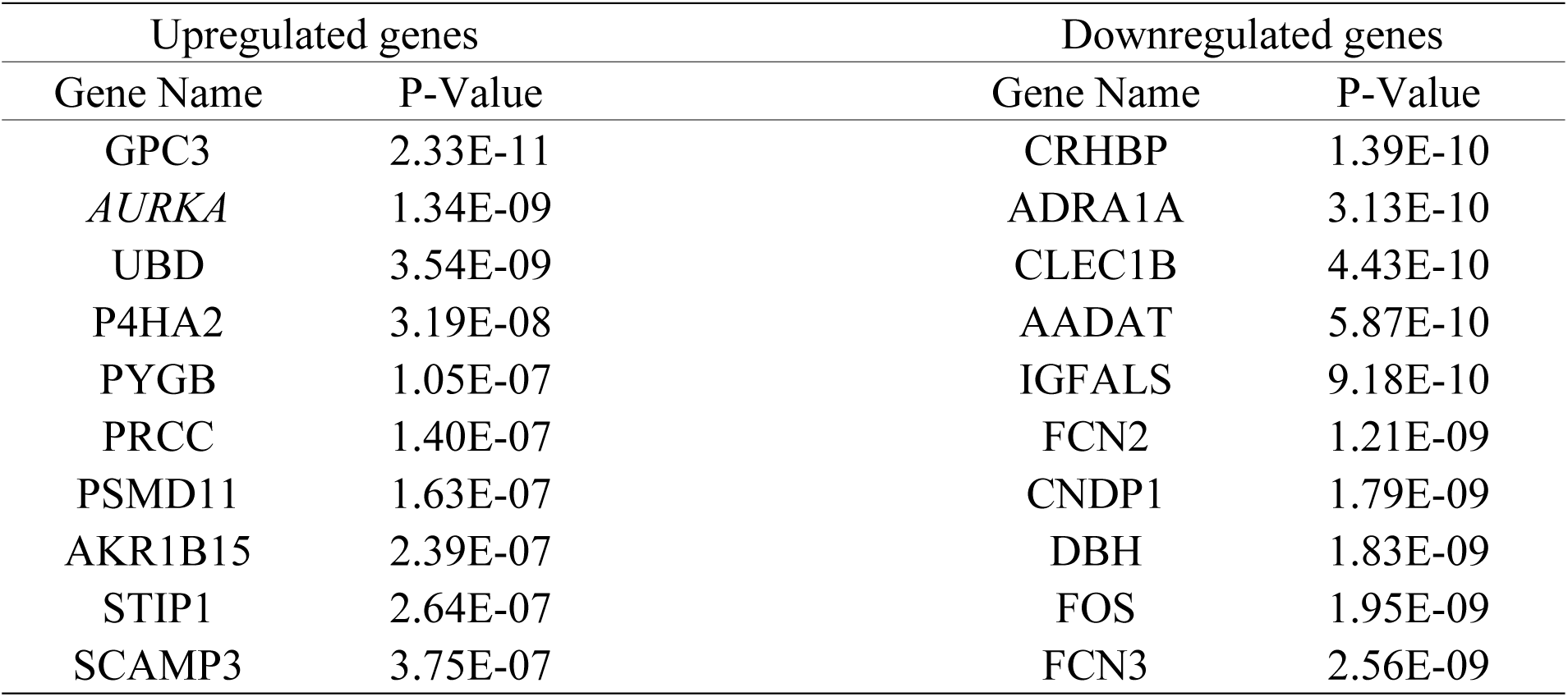
Top 10 most substantially up- or downregulated differentially expressed genes

| Upregulated genes |  | Downregulated genes |  |
| --- | --- | --- | --- |
| Gene Name | P-Value | Gene Name | P-Value |
| GPC3 | 2.33E-11 | CRHBP | 1.39E-10 |
| *AURKA* | 1.34E-09 | ADRA1A | 3.13E-10 |
| UBD | 3.54E-09 | CLEC1B | 4.43E-10 |
| P4HA2 | 3.19E-08 | AADAT | 5.87E-10 |
| PYGB | 1.05E-07 | IGFALS | 9.18E-10 |
| PRCC | 1.40E-07 | FCN2 | 1.21E-09 |
| PSMD11 | 1.63E-07 | CNDP1 | 1.79E-09 |
| AKR1B15 | 2.39E-07 | DBH | 1.83E-09 |
| STIP1 | 2.64E-07 | FOS | 1.95E-09 |
| SCAMP3 | 3.75E-07 | FCN3 | 2.56E-09 |

### 3.2 Functional enrichment analysis

Upregulated genesTo identify the pathways that had the most significant involvement with the genes identified, DEGs were submitted to DAVID for GO and KEGG pathway analysis. GO analysis revealed that, in biological process terms, the DEGs were mainly enriched in ‘oxidation-reduction process’, ‘cell division’, ‘epoxygenase P450 pathway’ and ‘regulation of complement activation’. In cell component terms, the DEGs were mainly enriched in ‘extracellular exosome’, ‘cytosol’, ‘extracellular space’ and ‘extracellular region’. In molecular function terms, the DEGs were mainly enriched in ‘arachidonic acid epoxygenase activity’, ‘retinol dehydrogenase activity’ and ‘alcohol dehydrogenase activity, zinc-dependent’ (Fig. 3). KEGG pathway analysis demonstrated that DEGs were significantly enriched in ‘Cell cycle’, ‘Metabolic pathways’, and ‘DNA replication’ (Fig. 4).

**Fig 3.**
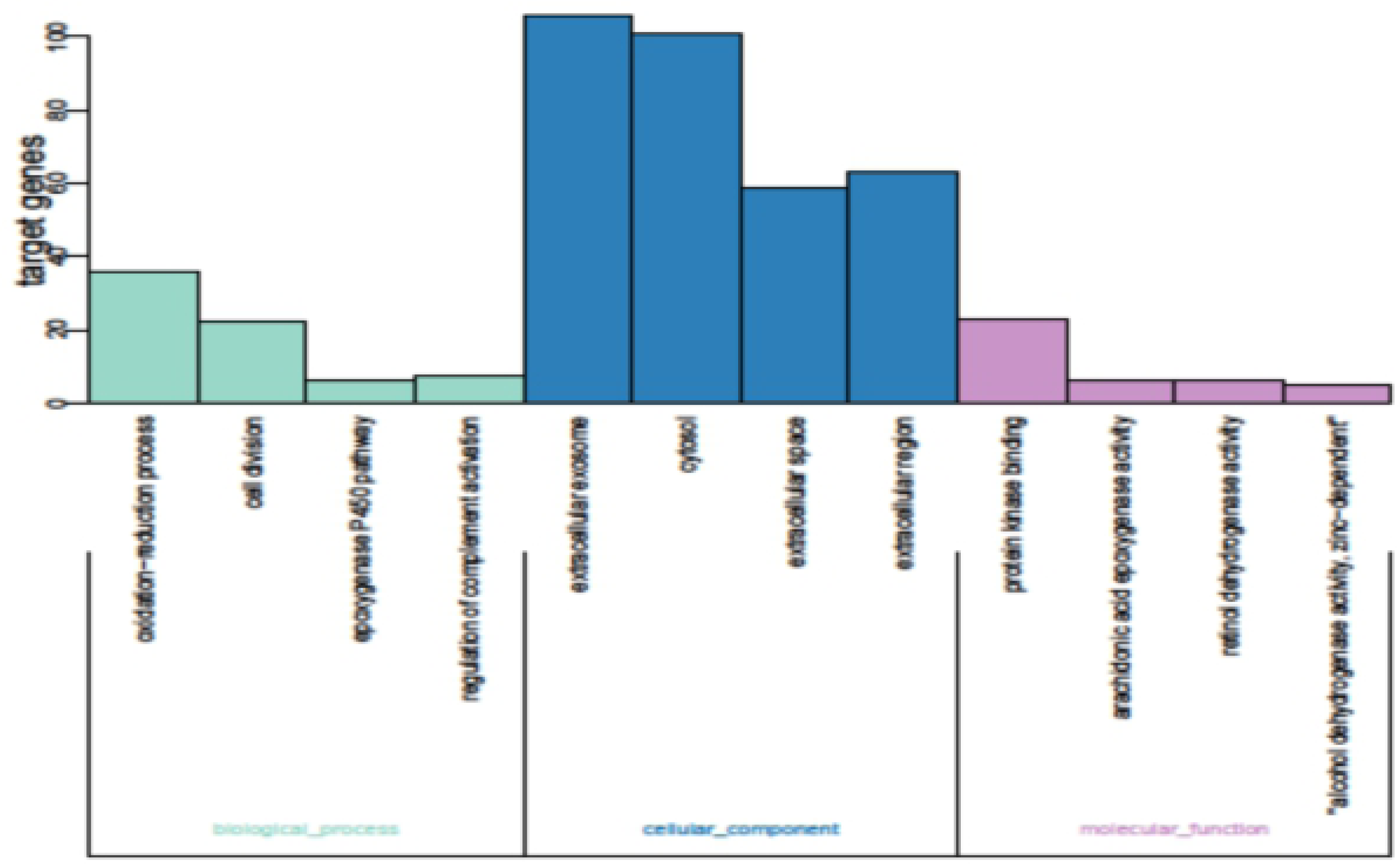
GO functional analysis of DEGs. GO enrichment analysis of DEGs was retrieved using DAVID. The 10 most significantly (P<0.05) enriched GO terms in biological process, molecular function and cellular component branches are presented. DEGs, differentially expressed genes; GO, gene ontology.

**Fig 4.**
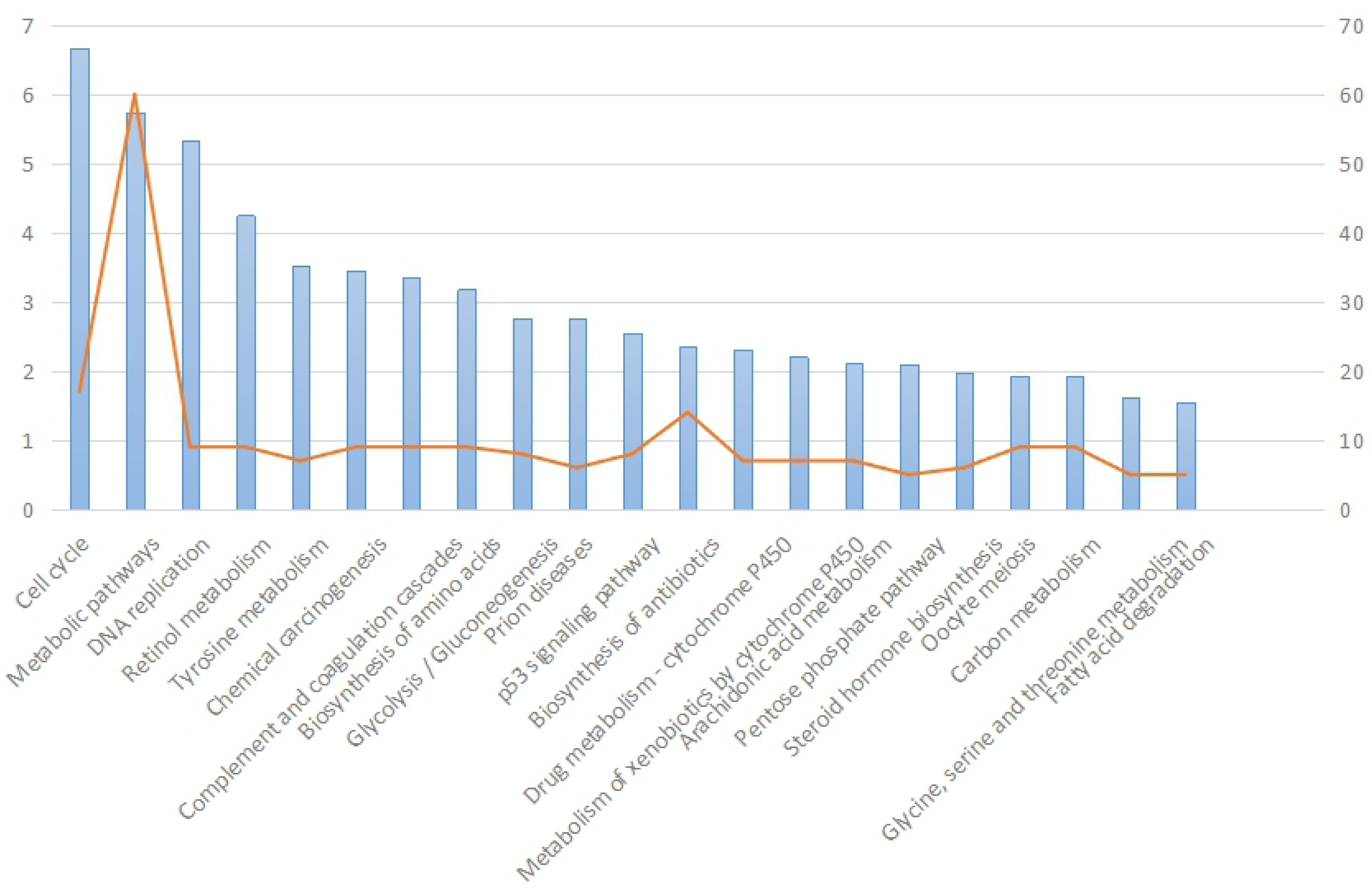
Significantly enriched pathways in KEGG pathway analysis of differentially expressed genes inhepatocellular. KEGG, Kyoto Encyclopedia of Genes and Genomes.

### 3.3 PPI network construction and module analysis

Interactions between the identified DEGs were revealed by constructing a PPI network (Fig. 5). In total, there were 366 nodes and 1440 edges in the network. According to degree levels, the top five hub nodes were cyclin A2 (*CCNA2*; degree, 25), polo like kinase 1 (*PLK1*; degree, 24), cell division cycle 20 (*CDC20*; degree, 18), ubiquitin conjugating enzyme E2 C (*UBE2C*; degree, 18) and kinetochore protein aurora kinase A (*AURKA*; degree, 17). A significant module was subsequently constructed with 32 nodes and 447 edges, which had the highest MCODE score (Fig. 6). Subsequent functional enrichment analysis revealed that the genes in this module were mainly enriched in ‘nucleoplasm’, ‘MCM complex’, ‘ATP binding’, ‘mitotic sister chromatid segregation’, and ‘DNA replication initiation’. GO and pathway analysis of genes in selected modules(Table 2).

**Fig 5.**
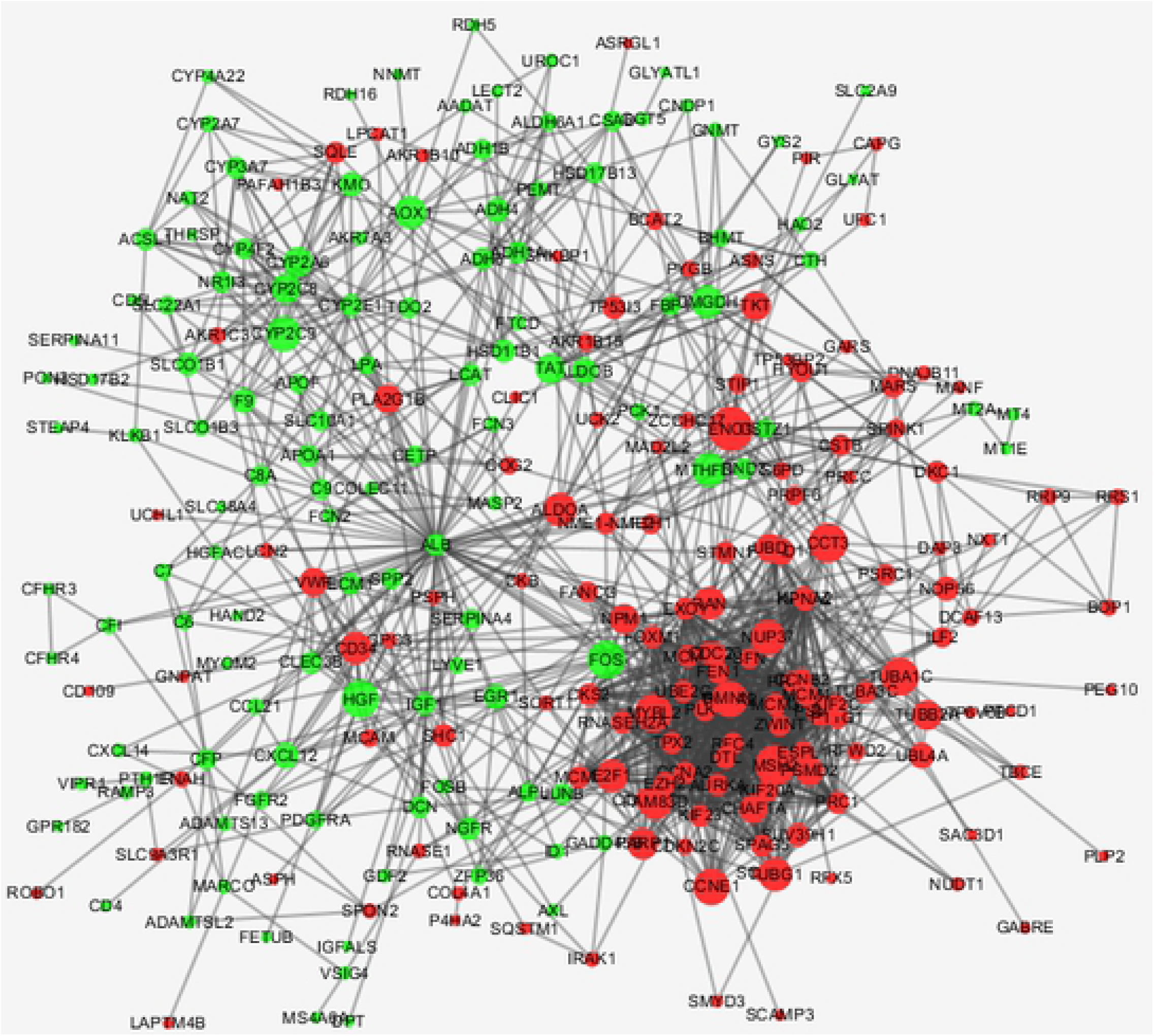
PPI network constructed from the common DEGs, module analysis and hub genes. Using the STRING online database, 366 DEGs were filtered into the DEG PPI network complex. The nodes represent proteins, the edges represent the interaction of proteins and green circles and the red circles indicate downregulated and upregulated DEGs, respectively. PPI, protein-protein interaction; DEG, differentially expressed gene.

**Fig 6.**
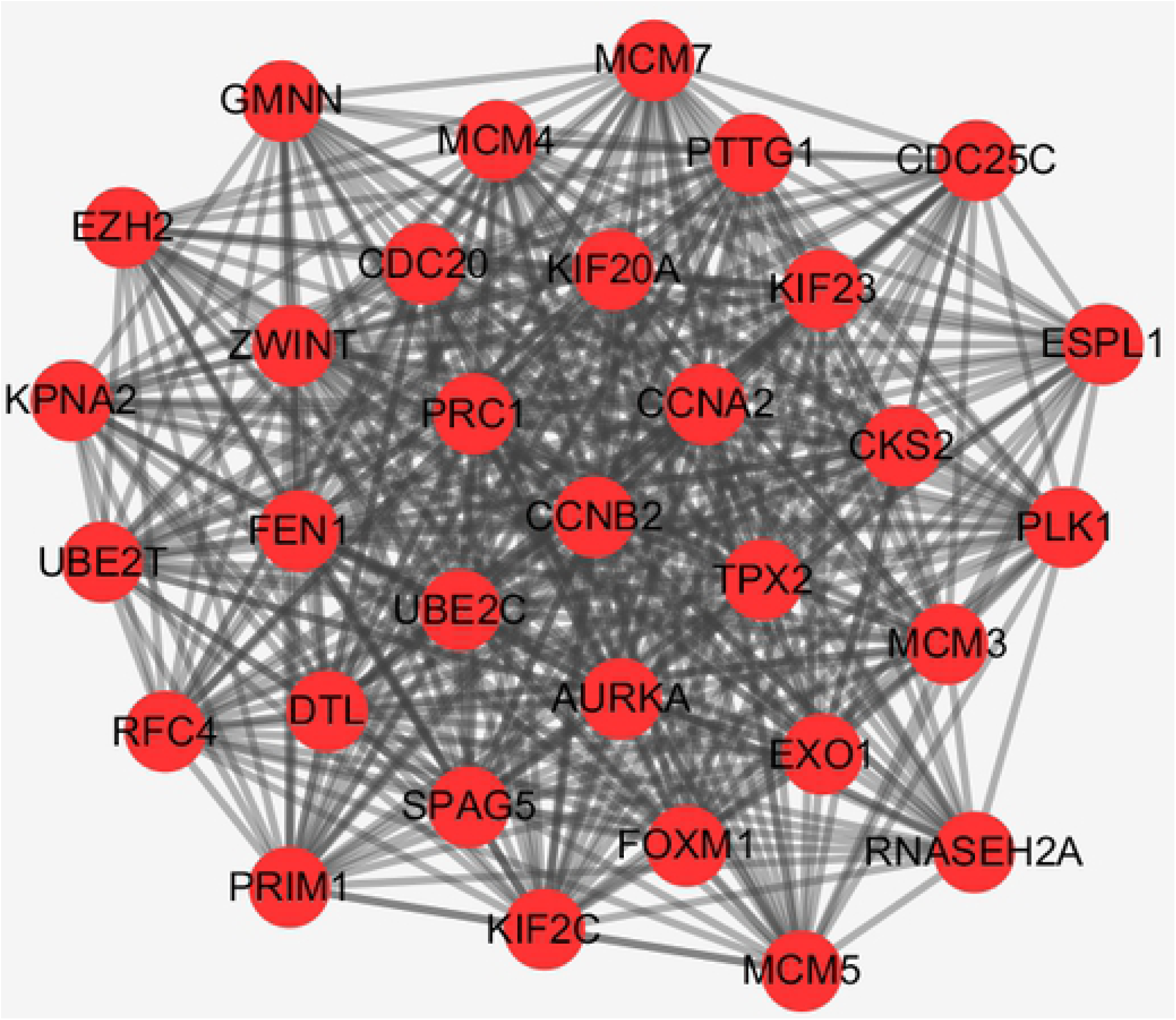
The highest scored module in the PPI network. PPI, protein-protein interaction.

**Table 2.** GO and pathway analysis of genes in select modules

| Category | Pathway ID | Term | Count | PValue |
| --- | --- | --- | --- | --- |
| BP | GO:0000070 | mitotic sister chromatid segregation | 4 | 9.34E-06 |
| BP | GO:0006270 | DNA replication initiation | 4 | 1.52E-05 |
| BP | GO:0045143 | homologous chromosome segregation | 3 | 1.60E-05 |
| CC | GO:0005654 | nucleoplasm | 15 | 8.64E-08 |
| CC | GO:0042555 | MCM complex | 4 | 5.02E-07 |
| CC | GO:0005737 | cytoplasm | 17 | 1.70E-05 |
| CC | GO:0000784 | nuclear chromosome, telomeric region | 5 | 3.45E-05 |
| MF | GO:0005524 | ATP binding | 12 | 6.09E-06 |
| KEGG | cfa04110 | cell cycle | 11 | 1.43E-12 |
| KEGG | cfa03030 | DNA replication | 8 | 1.03E-11 |
| KEGG | cfa04114 | oocyte meiosis | 6 | 1.89E-05 |

### 3.4 Validation of target protein mRNA levels in HCC

Based on TCGA, it was identified that the RNA expression levels of ‘*CCNA2*’, ‘*PLK1*’, ‘*CDC20*’, ‘*UBE2C*’ and ‘*AURKA*’ were significantly increased in HCC samples as compared to normal liver samples(Fig. 7).

**Fig 7.**
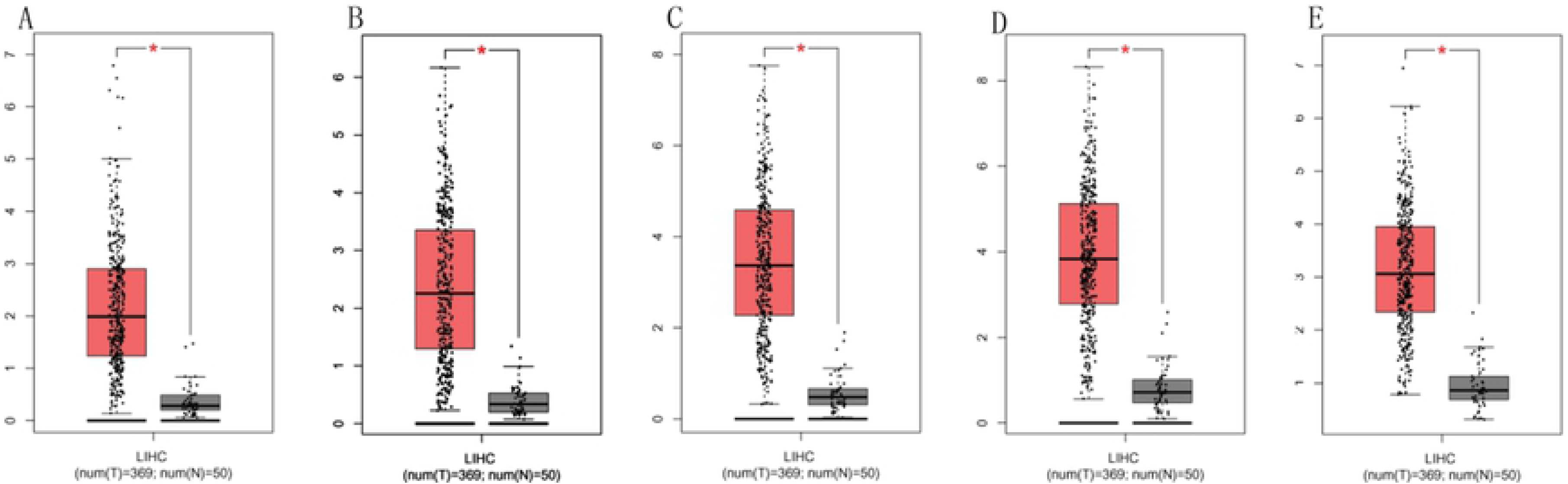
Analysis of the expression of hub genes in TCGA. RNA expression levels of (A) CCNA2, (B) PLK1, (C) CDC20, (D) UBE2C and (E) AURKA in normal liver vs. HCC compared.

We performed univariate Cox regression analysis. The results showed that the five hub genes were significantly correlated with the overall survival(log-rank P <.05) and disease free survival (logarithmic rank P <.05) of HCC patients (Fig. 8). These findings suggest that these hub genes may be a useful candidate for survival prediction in HCC patients.

**Fig 8.**
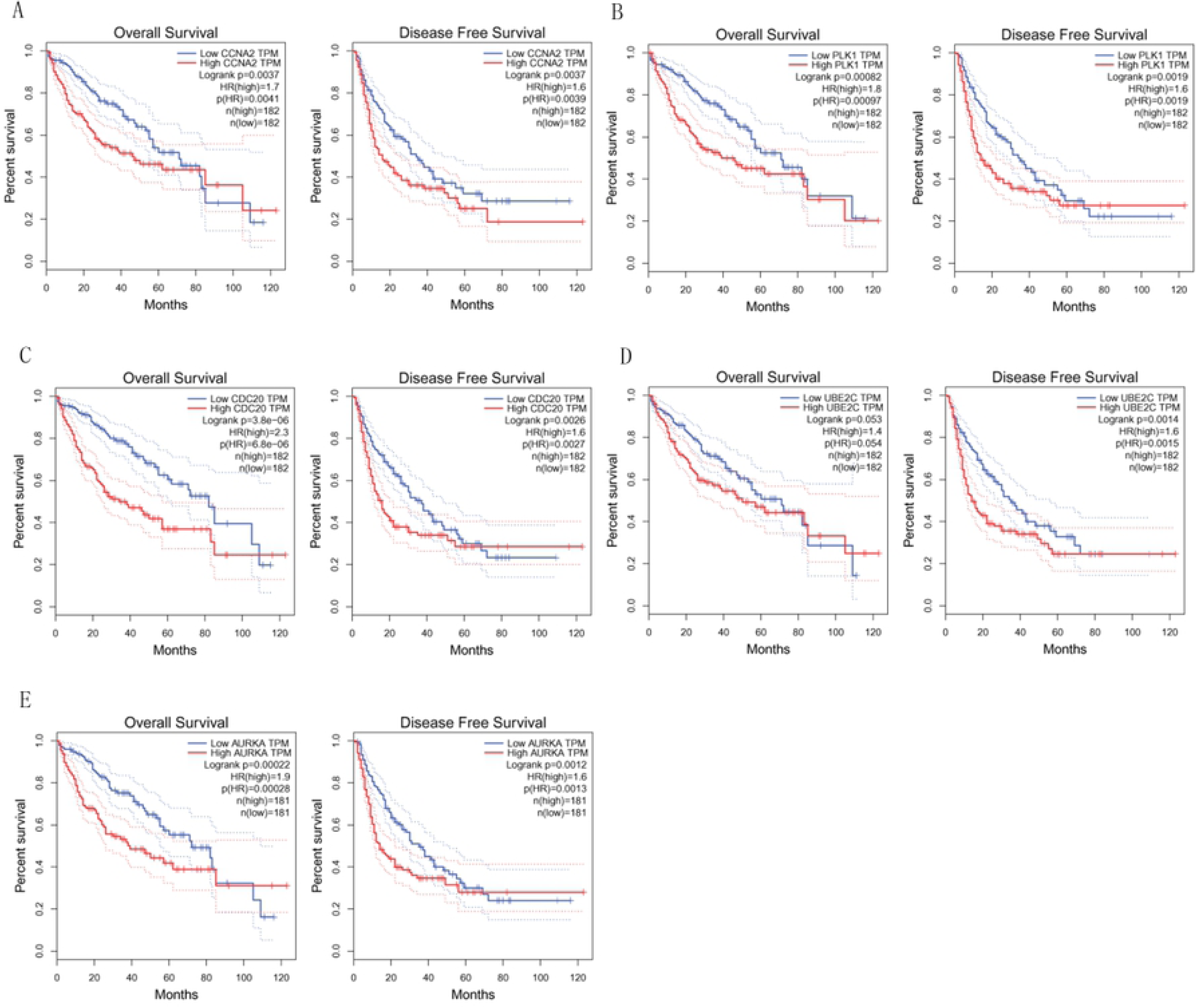
The Kaplan-Meier curves the OS and RFS of HCC patients in the TCGA. (A) CCNA2, (B) PLK1, (C) CDC20, (D) UBE2C and (E) AURKA in normal liver vs. HCC were compared.

### 3.5 Upstream miRNA analysis

According to the information in starBase v2.0, the miRanda prediction program was selected to search for miRNAs upstream of the five hub genes. By comparing DEM targets, *CCNA2* was found to be a potential target of 48 miRNAs, *AURKA* was found to be a potential target of 31 miRNAs, *UBE2C* was found to be a potential target of nine miRNAs, and *PLK1* was found to be a potential target of three miRNAs. *CDC20* was not found to be a potential target of miRNA. Notably, upstream miRNA analysis showed that *mir-300* and mir-381-3p were common upstream miRNAs of *CCNA2*, *UBE2C* and *AURKA*

## 4. DISSCUSION

HCC is one of the most common malignant tumors in the clinic, and the incidence rate is third among all malignant tumors[14]. Although the research on HCC has made great progress in recent years [15], its pathogenesis is not yet fully understood. Due to the biological diversity and complexity of individuals, the early diagnosis and treatment of HCC remains enigmatic. With the rapid development of high-throughput sequencing technology, an increasing number of gene functions have been recognized, and the early diagnosis and treatment of HCC can be enhanced by identifying target genes and related pathways[16].

In this study, we analyzed the GSE84004 and GSE54283 gene chips, extracted relevant differential genes, and analyzed 359 DEGs. To explain the common biological functions of these DEGs, we performed GO and KEGG analysis on the DAVID platform. In terms of biological process, prior studies have reported that the redox process is essential for maintaining cell homeostasis, which can maintain cell proliferation[17]. Disruption of the redox process causes oxidative damage to lipids, proteins and DNA, and thereby promotes tumor cell growth. In addition, the epoxygenase cytochrome P450 pathway plays a central role in various cellular functions. At the same time, cell division and regulation of complement activation plays an onocogenic role during tumor initiation and progression[18].

In terms of cell composition, the metabolism of most substances in the body and the modification of some proteins are carried out in the cytosol, and the unlimited growth potential of tumor cells depends on amino acid availability in the cytosol as well as metabolic energy[19]. Recently, researchers have found that carcinogenic exosomes can facilitate immune escape, and when carcinogenic exosomes exceed tumor suppressive signaling, they can eventually cause tumorigenicity in the microenvironment[20, 21]. In terms of molecular function, arachidonic acid cyclooxidase, retinol dehydrogenase and alcohol dehydrogenase are involved in the production of cytochrome P450, retinoic acid and other factors, respectively. These factors play an important role in the metabolism of many biologically important substances, catalyzing oxidation or reducing the specificity of multiple substrates, and maintaining the activities of tumor cells[22]. In addition, zinc staining in liver tissues confirmed that zinc is significantly lower in HCC cells than in untransformed liver cells[23]. Analysis of the KEGG pathway revealed that these DEGs were mainly enriched in ‘Cell cycle’, ‘Metabolic pathways’ and ‘DNA replication’[24–26]. To predict the associations and protein functions of these DEGs, a PPI network was constructed in which the top five central genes with the highest degrees of connectivity were selected, including *CCNA2*, *PLK1*, *CDC20*, *UBE2C* and *AURKA*. Tumor development is an extremely complex process in which many genetic and epigenetic modifications of cancer-driving genes occurs. Module analysis of the PPI network also shows that these genes are associated with ‘cell cycle’, ‘DNA replication’ and ‘Oocyte meiosis’ signaling pathways. In addition, to verify the expression levels of these central genes, we obtained clinical information and RNA sequencing data of HCC samples through TCGA and conducted an analysis. We found that the five hub genes showed significant differences in tumor and corresponding non-tumor tissues. Then, we performed univariate Cox regression analysis. The results showed that the five hub genes were significantly correlated with the total survival time and the survival time without recurrence of HCC patients.

Cyclin A2, the protein encoded by the *CCNA2*gene belongs to the highly conserved cyclin family whose members function as regulators of the cell cycle. This protein binds to and activates cyclin-dependent kinase 2 and thus promotes the G1/S and G2/M transitions[27]. Many studies have shown that genetic alterations in *CCNA2* have been identified in several types of malignancies, such as esophageal cancer breast cancer [28, 29]. It has recently been demonstrated that *CCNA2* interacts with actin and RhoA during mitosis and induces a significant decrease in active RhoA in mitosis by depletion, thereby promoting cell proliferation[30].

Polo like kinase 1, the Ser/Thr protein kinase encoded by the *PLK1*gene, belongs to the CDC5/Polo subfamily. It is highly expressed during mitosis, and elevated levels are found in many different types of cancer. Depletion of this protein in cancer cells dramatically inhibits cell proliferation and induces apoptosis; hence, it is a target for cancer therapy. *PLK1* coordinates biosynthesis during cell cycle progression by directly activating the pentose phosphate pathway and promotes RNF2 degradation to regulate cell mitosis to ultimately regulate tumorigenesis [31, 32].

*CDC20* encodes cell division cycle 20, which is a regulator of cell cycle checkpoints. It binds directly to another regulatory factor, Cdh1, and activates the late mitotic promoting complex APC, which plays an important role in late-stage of cell division and withdrawal from mitosis [33, 34]. Recently, an increasing number of studies have shown that *CDC20* is a carcinogenic factor that is widely regulated in a variety of human malignancies, including lung cancer, kidney cancer and prostate cancer. Furthermore, *CDC20* is closely related topoor prognosis of a tumor [35–37].

*UBE2C* encodes ubiquitin conjugating enzyme E2 C, a member of the E2 ubiquitin conjugating enzyme family, which includes ubiquitin-activating enzymes, ubiquitin-conjugating enzymes and ubiquitin-protein ligases[38], which collectively tag proteins for proteasomal degradation, by which mitotic cyclins are destroyed and cell cycle progression is promoted[39]. *UBE2C* plays a role in a variety of cancers, including Esophageal cancer, gastric cancer and lung cancer [40–42].

*AURKA* encodes aurora kinase A which, as a member of the Aurora kinase family[43], regulates the G2/M transition during mitosis by regulating thespindle, centrosomes and chromosome segregation [44–46]. Prior studies have shown that AURKB overexpression can lead to multinuclear cells and polyploidy [47] and is highly correlated with genomic instability and poor prognosis in neuroblastoma [48]. Other studies have shown that *AURKA* can promote the expression of anti-apoptotic proteins in tumor cells, and this gene can be used as a biomarker and target for a variety of tumor treatments [49–51].

In this study, we found that all five hub genes were related to the cell cycle. The results of miRNA analysis showed that *CCNA2*, *AURKA* and *UBE2C* were potential target genes shared by *mir-300* and *mir-381-3p*. Prior studies have shown that the expression of *mir-300* and *mir-381-3p* in gastric cancer, thyroid cancer, osteosarcoma and other tumors is disordered[52–55]. Although the mechanisms of *mir-300* and *mir-381-3p* in HCC are not clear, we hypothesize that *mir-300* and *mir-381-3p* may alter the cell cycle by regulating *CCNA2*, *AURKA* and *UBE2C*, thereby promoting the proliferation of liver cancer cells.

In summary, this study identified candidate genes and pathways that may be involved in HCC progression using a comprehensive analysis of two cohort data sets. These results may facilitate a deeper understanding of the molecular mechanisms underlying HCC and provide a range of potential biomarkers for future research. These findings add important insights into the diagnosis and treatment of HCC. However, the lack of experimental validation is a limitation of our research. Therefore, further experimental research is needed to verify these findings with a larger sample size.

## 6. Supporting information

**S1 Table. DEGs screened by the GSE84040 and GSE54238 data sets.**

**S2 Table. Gene ontology analysis of the DEGs.**

**S3 Table. KEGG pathway enrichment analysis of the DEGs.**

**S4 Table. PPI network nodes of DEGs analyzed using Cytoscape software.**

